# Triplex real-time PCR ZKIR-T assay for simultaneous detection of the *Klebsiella pneumoniae* species complex and identification of *K. pneumoniae* sensu stricto

**DOI:** 10.1101/2024.02.02.578617

**Authors:** Małgorzata Ligowska-Marzęta, Elodie Barbier, Carla Rodrigues, Pascal Piveteau, Dennis Schrøder Hansen, Alain Hartmann, Eva Møller Nielsen, Sylvain Brisse

## Abstract

*Klebsiella pneumoniae* species complex (KpSC) members, including the most important species *K. pneumoniae* (phylogroup Kp1 of the KpSC), are important opportunistic pathogens which display increasing rates of antimicrobial resistance worldwide. As they are widespread in food and the environment, there is a need for fast, sensitive and reliable methods to detect KpSC members in complex matrices. Previously, the ZKIR real-time PCR assay was developed to detect all KpSC members without distinction. Given that Kp1 is the clinically most significant phylogroup of the KpSC, here we aimed to simultaneously identify Kp1 while detecting all KpSC members. Three TaqMan probes were developed and used: the *zkir* P1 probe to detect phylogroups Kp1 to Kp5 and Kp7; the *zkir* P2 probe to detect phylogroup Kp6; and the Kp1 probe to specifically identify this phylogroup. The new assay was tested on a total of 95 KpSC and 19 non-KpSC strains from various sources, representing the different phylogroups as defined by whole genome sequencing. The results showed almost complete specificity, as the expected PCR results were obtained for 112 (98%) strains. The new triplex real-time PCR assay, called ZKIR-T, enables detection of all KpSC taxa while discriminating Kp1, which will be useful for rapid screening and to focus downstream analyses on chosen phylogroups of the KpSC.

**Importance:** The pathogens of the *Klebsiella pneumoniae* species complex are widespread in food and animals and are amongst the main pathogens responsible for multidrug resistant infections in humans. In this study, we developed a highly sensitive detection assay that enables detection of this group of bacteria, with the simultaneous identification of the most common and clinically important species. This triplex one-reaction assay was shown to be highly sensitive and precise, enabling fast screening of varied samples for the presence of KpSC and *K. pneumoniae sensu stricto*.

## Introduction

The *Klebsiella pneumoniae* species complex (KpSC) comprises five different species organized into seven distinct phylogroups (1, 2): Kp1 (*K. pneumoniae sensu stricto*), Kp2 (*K. quasipneumoniae* subsp. *quasipneumoniae*), Kp3 (*K. variicola* subsp. *variicola*), Kp4 (*K. quasipneumoniae* subsp. *similipneumoniae*), Kp5 (*K. variicola* subsp. *tropica*), Kp6 (‘*K. quasivariicola’*) and Kp7 (*K. africana*) (2–4). Among them, phylogroup Kp1 is the most frequently found in clinical settings, accounting for ∼85% of the isolates identified as *K. pneumoniae* by classical microbiological methods (e.g. MALDI-TOF MS) (1).

*K. pneumoniae*, which is a member of the ESKAPE pathogen group (*Enterococcus faecium, Staphylococcus aureus, Klebsiella pneumoniae, Acinetobacter baumannii, Pseudomonas aeruginosa* and *Enterobacter* species) (5) poses a serious infectious risk, especially due to occurrence of multidrug-resistant (MDR) *K. pneumoniae*, such as extended-spectrum β-lactamase (ESBL) and/or carbapenemase producers (1, 6). Even though *K. pneumoniae* is dominant in clinical settings, the other members of the KpSC also cause infections and may be more frequent in other habitats. Therefore, investigating the distribution of KpSC members in other ecological niches is important to improve knowledge on their ecology and transmission (7). Members of the KpSC complex have been isolated from a variety of sources, such as dust samples from pig holdings, air samples in broiler houses, dairy cows milk, retail meat, seafood, vegetables, birds, insects and other mammals (8–10). High prevalence of KpSC (∼50%) was also found in chicken meat and salads, with a predominance of the Kp1 (91%) and Kp3 (6%) phylogroups (11). A comparative study of human and bovine isolates showed that the Kp3 phylogroup was dominant among bovine isolates, in contrast to Kp1 and Kp2 dominating among human isolates (12). *KpSC* members are also found in water, plants and soil (13). A large analysis reported on the wide diversity of *Klebsiella* isolates in the environment (14). Samples collected from clinical, community, veterinary, agricultural and environmental sources revealed that half of the isolates (N=1705) were *K. pneumoniae* and that this species’ primary sources were hospitals and livestock. The other members of the KpSC were found as well: *K. variicola* (N=279), *K. quasipneumoniae* subsp. *quasipneumoniae* (N=76), *K. quasipneumoniae* subsp. *similipneumoniae* (N=49) and *K. quasivariicola* (N=4).

The ubiquity of KpSC bacteria in the environment and their importance in clinical settings underline the need to better understand their ecology and transmission. Reliable and effective methods for their detection and identification from a variety of samples are therefore essential. We previously described a real-time PCR method, called the ZKIR assay, with the aim of detecting KpSC isolates from complex matrices such as soil samples (15). The method has since been used on other complex matrices such as food (11) and human gut samples (16), where its sensitivity and accuracy have been validated against traditional culturing methods (SCAI) and whole metagenomic sequencing (WMS). However, the ZKIR assay is unable to discriminate strains between the individual KpSC phylogroups. The aim of this study was to address this limitation by developing and validating a real-time PCR assay that can detect all KpSC phylogroups and simultaneously identify its most important member, Kp1 (*K. pneumoniae sensu stricto*, or *K. pneumoniae* for short).

## Results

### Probes and primers design

To adapt the SYBR Green ZKIR method (15) to a Taqman PCR assay, a ZKIR probe, called ZKIR_P1, was designed using Primer3Plus (https://www.bioinformatics.nl/cgi-bin/primer3plus/primer3plus.cgi). The specificity of the primers, the predicted probe and the amplicons was confirmed by applying BLASTN on the GenBank nucleotide collection (nr/nt) of Kp1 to Kp5 genomes from the NCBI database (no Kp6 genome was available). The *in silico* alignment of the ZKIR_P1 probe against six available Kp6 genomic sequences provided by Institut Pasteur showed numerous single nucleotide polymorphism (SNPs), which could potentially prevent hybridization with the Kp6 target sequence. The qPCR assays using ZKIR primers and P1 probe were carried out to test this hypothesis and confirmed the lack of amplification (data not shown). Therefore, a Kp6 specific probe, called ZKIR_P2, was additionally designed for Kp6 amplification (see Table 1).

**Table 1.**
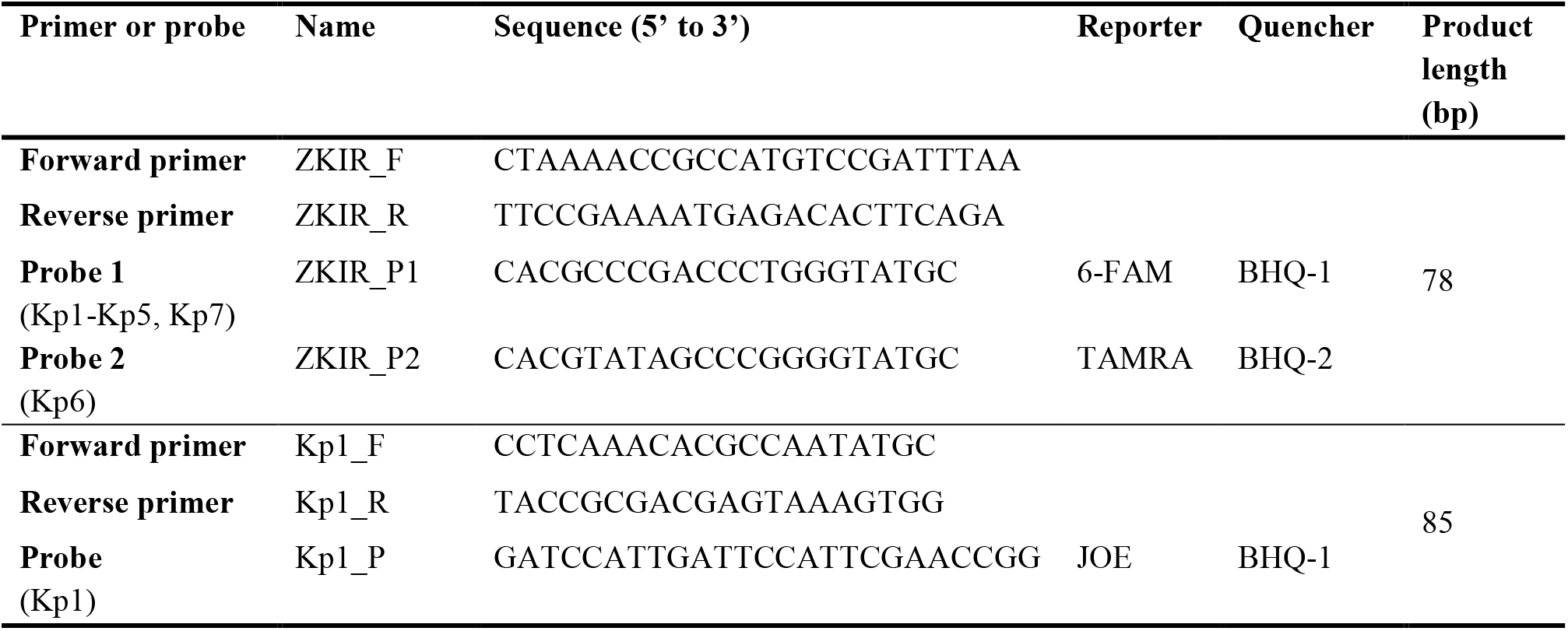
Primers and probes developed and used in this study.

Four molecular targets have previously been proposed for Kp1 specific detection, and were assessed *in silico* for their specificity (15). The conclusion was that these targets were not specific for Kp1, as they were also detected in other KpSC phylogroups and/or in *K. aerogenes* and *K. oxytoca* (see Figure S1 in Barbier *et al*, 2020). We therefore conducted a pangenome mapping and comparative genomics strategy to identify specific targets for the different KpSC phylogroups, including Kp1. As a result, a Kp1-specific and conserved gene was selected. This 843-bp gene is annotated as coding for the helix-turn-helix transcriptional activator NimR, a transcriptional activator of the efflux pump of the MFS family, NimT (for nitroimidazole transporter), responsible for the 2-nitroimidazole resistance in *E. coli* (17). The specificity of this Kp1-specific gene candidate was checked using BLASTN with customized parameters, showing that *nimR* was only detectable in Kp1 phylogroup genomes.

Several sets of primers and probes were then designed and their *in silico* specificity investigated. One set, called Kp1, was fully specific for Kp1 phylogroup and was expected to amplify an 85-bp DNA fragment (Table 1).

The ZKIR_P1_P2 duplex and ZKIR_P1_P2/Kp1_P (ZKIR-T) triplex real time PCR assays were separately tested on 49 KpSC strains and 19 non-KpSC isolates (Table S1). Screening with the duplex system showed that it amplified specifically all the members of the KpSC (Kp1 to Kp5 and Kp7) except Kp6 with the ZKIR_P1 probe, whereas the Kp6 strains were amplified with the ZKIR_P2 probe. Regarding the triplex assay, results were in complete agreement with the duplex ones with equivalent cycle thresholds (Ct) (data not shown). The 9 Kp1 isolates were correctly detected in the triplex system (Table S1), whereas no non-KpSC isolates were amplified with any of the probes.

### Validation of the triplex real time PCR assay on a distinct KpSC collection

The method was then implemented in a second laboratory and tested using local samples. Forty-six additional KpSC strains, previously sequenced by WGS, were tested at Statens Serum Institut (SSI) with the triplex assay. The Kp1_P system used in the real time PCR was found to be specific towards Kp1 phylogroup; the ZKIR_P1 system towards phylogroups Kp1-Kp5 and Kp7, and the ZKIR_P2 system towards Kp6 isolates. However, there were two exceptions, leading to specificity of 96% (44 out of 46 test strains), when compared with species assignation from genomic data. The results, together with the Ct values for each Kp phylogroup can be seen in Table S3.

One *K. variicola* subsp. *variicola* (Kp3) isolate (MVK-06S035) was detected by the ZKIR_P1 probe, as expected, but at a higher Ct than that of the Kp3 reference strain and only in 1 out of 3 replicates. Sequence alignment with the ZKIR primers and the ZKIR_P1 probe of the genomic sequence of this isolate revealed mismatches in the forward primer (3 nt) and the probe region (4 nt) (Figure S 1A), which is a likely explanation for the late amplification and lack of reproducibility between the replicates. Another Kp3 isolate (MVK-06H168) was detected by the ZKIR_P1 probe in all three replicates, but also with higher Ct values than for the Kp3 reference strain. Alignments of ZKIR primers and P1 probe of its genomic sequence also showed mismatches, mainly in the forward primer (4 nt) and the probe region (2 nt) (Figure S 1B).

## Discussion

Here, we addressed a so-far unmet need for surveillance and research studies on the ecology and transmission of KpSC isolates. Whereas the previously existing real-time ZKIR PCR assay allowed for the detection of KpSC members as a whole (15), it could not distinguish between individual phylogroups.

We thus developed an extension of the previous tool, able to detect phylogroup Kp1 (*K. pneumoniae*) as well as the members of the KpSC. The novel triplex real-time PCR assay developed herein will enable screening of samples for the presence of Kp1, the clinically most concerning member of the complex, or to detect samples where the other species, which are less abundant, may be present and subsequently analyzed, *e*.*g*., by culture and downstream characterization. The probe ZKIR_P1, designed in this study for the development of a multiplex assay, detected all phylogroups except Kp6 and was therefore combined with a second probe, ZKIR_P2, which is Kp6-specific, in order to consistently detect all KpSC phylogroups. This adaptation to include Kp6 was not necessary in the initial ZKIR assay, as Kp6 is appropriately detected using this SYBR green assay due to the absence of mismatch on the priming sites. The novel Kp1_P system targets a transcriptional regulator NimR and was designed to be specific for the Kp1 phylogroup based on a large genomic dataset. These Kp1-specific primers and probe (the Kp1_P system) were used in simplex qPCR in another study (18) for identification of human, animal and environmental isolates (N= 433). Assignments of Kp1 (n = 256/433; 56.6 %) were 100% concordant with the whole-genome sequencing (WGS) results. No other phylogroup member was amplified with this set of Kp1 PCR primers and probe.

Hence, the resulting triplex real time PCR, which we named the ZKIR-T assay (“T” for three) now allows simultaneous detection of all members of KpSC and a distinction on whether the isolate belongs to Kp1, Kp6 or one of the other phylogroups.

The new triplex real time PCR assay was tested on a total of 114 isolates in two different laboratories and led to identification consistent with the WGS results in one reaction with 98% specificity. Mismatches of the forward primer and probe regions with the target region of two *K. variicola* (Kp3) isolates were responsible for the lack of amplification of the ZKIR target in one isolate and for the amplification at a higher Ct value in the other isolate. Despite the two test strains that showed unexpected results, we consider the method was validated and could be recommended,e.g. for studies on the occurrence of KpSC and simultaneous identification of Kp1 in large sample collections, as was done previously for the ZKIR qPCR to screen food samples for the presence of KpSC members. Still, studies of larger sample or isolates sets are needed to further evaluate the accuracy, analytical sensitivity and specificity of the ZKIR-T real time PCR assay.

## Acknowledgments

We thank Ibado Mahad (SSI) for expert technical assistance in the laboratory and Arnaud Dechesne (Technical University of Denmark – DTU) and the staff at VandCenter Syd, Odense, for providing sewage samples.

S.B., E.M.N. and P.P. conceptualized the study. E.B., A.H. and C.R. designed the primers and probes and performed validation of the ZKIR-T assay. D.S.H. isolated strains from human clinical sources.

M.L.M. performed the ZKIR-T assay on another set of samples. C.R. and M.L.M. analysed the genomic data. M.L.M., E.B., C.R. and S.B. wrote the original draft of the manuscript. All authors contributed to writing and editing the manuscript and reviewed the final version.

## Open access

This research was funded, in whole or in part, by Institut Pasteur and by European Union’s Horizon 2020 research and innovation programme. For the purpose of open access, the authors have applied a CC-BY public copyright license to any Author Manuscript version arising from this submission.

## Materials and methods

### *In silico* analyses for target selection

The ZKIR region was previously proposed as target for the specific detection of KpSC members in a SYBR green PCR assay (15). The pair of primers (ZKIR_F and ZKIR_R) specifically amplifies a 78-bp sequence in the intergenic region (IR) located between *zur* (zinc uptake regulator) and *khe* (annotated as a putative haemolysin) genes. The specificity and the sensitivity of this system was already demonstrated on a wide variety of samples (11)(18)(16). With the objective to develop a multiplex assay, a ZKIR probe (ZKIR_P1) was designed using a multiple alignment tool (SeaView) and primers and probes using design interface (Primer3Plus). A second probe (ZKIR_P2) was also designed against six available Kp6 genomic sequences provided by Institut Pasteur. This probe targets the same locus as ZKIR_P1 but differs in 6 nucleotides.

The design of a Kp1 specific system was recently described in another study (Supplemental File 1 from Dereeper et al., 2022). Briefly, a collection of 66 reference genomes from Institut Pasteur, including representatives of Kp1 to Kp6 phylogroups, was used to construct a pangenome with Roary v3.11 (19) using a blastP identity cut-off of 80% and core genes defined as those being present in more than 90% of the isolates. From the pangenome, we have selected genes exclusive of each phylogroup (i.e. unique genes for Kp1, Kp2, etc), with a total of 8 candidate target genes being obtained for Kp1. The specificity of each candidate was checked using BLASTN with customized parameters against: (i) all available *Klebsiella* spp. genomes; and (ii) all available genomes excluding *Klebsiella* spp. (last accessed: February 2018). The phylogenetic distribution of the Kp1 candidate target gene retained (*nimR*) was assessed using BLASTN (80% nucleotide identity and 80% length coverage as cutoffs) in a previous genomic collection of 1,001 genomes from NCBI (February 2018) and Institut Pasteur’s internal collection, which included *Klebsiella* spp. and closely related species (*Raoultella* spp.) that were mapped in a phylogenetic tree.

### Validation of the ZKIR_P1_P2/Kp1_P triplex assay (ZKIR-T) using an in-house collection

Two assays were carried out in parallel using ZKIR_P1_P2 duplex system for the first one and ZKIR_P1_P2/Kp1_P triplex system (ZKIR-T) (Table 1) for the second on a reference dataset panel from INRAe, which included 49 KpSC strains and 19 closely related species (Table S 1). In addition to that, eight control strains representing each of the 7 phylogroups (Table S 2) and 46 KpSC test strains representing phylogroups Kp1 to Kp6, without the Kp5 group, (Table S 3) were used in validation of the triplex PCR at SSI. Hence, in total, 95 KpSC and 19 non-KpSC strains were used for the ZKIR-T aassay validation. This collection was identified based on genomic sequences and their distance to type strains of the seven taxa of the KpSC. It consisted of 20 *K. pneumoniae* (Kp1), one *K. quasipneumoniae* subsp. *quasipneumoniae* (Kp2), 20 *K. variicola* (Kp3), 4 *K. quasipneumoniae* subsp. *similipneumoniae* (Kp4) and one *K. quasivariicola* (Kp6). The isolates were selected from larger collections from various sources (Table S 3), such as sewage, food, animal carriage, human faeces and clinical blood samples, in order to represent as many of the 6 phylogroups from the complex as possible. Primers and probes presented in Table 1 were used for validation.

### DNA extraction method for real-time PCR

The bacterial DNA of the reference strains was extracted at INRAe (France) according to an in-house protocol based on a phenol/chloroform method as described in Barbier et al., 2020. At SSI (Denmark), the DNA was purified using DNeasy Blood and Tissue kit (Qiagen) following the standard protocol. Concentrations were measured using Qubit fluorometer (Invitrogen, Waltham, MA, USA).

### Real-time PCR

Multiplex PCRs were carried out using the primers and probes listed in Table 1. Reactions were carried out in a 20 µL reaction mix containing 10 µL of Takyon ROX Probe MasterMix Blue dTTP (Eurogentec) at a final concentration of 1X, 1 µL of each primer (final concentration 500 nmol l^-1^), 1 µL of each probe (final concentration 180 nmol l^-1^), 0.5 µL of ultrapure water and 2.5 µL of DNA.

Real-time PCR assays were performed in a StepOnePlus Real-Time PCR System (Thermo Fischer Scientific) at INRAe (France) and in a QuantStudio 5 Real-Time PCR System (Thermo Fischer Scientific) at SSI (Denmark). Initial denaturation was performed at 95°C for 3 min, followed by 40 cycles with denaturation at 95°C for 15 s and annealing and elongation at 60 °C for 1 min, as described in protocols.io: dx.doi.org/10.17504/protocols.io.b4s5qwg6. The specificity and the cross reactivity of the assays were evaluated at INRAe with purified DNA (1.8 to 4.4 ng per µL depending on the strains) of 49 reference KpSC members comprising 9 Kp1, 9 Kp2, 9 Kp3, 9 Kp4, 7 Kp5, 5 Kp6 and 1 Kp7 and 19 non-KpSC isolates, belonging to closely related species such as *Klebsiella aerogenes, Raoultella* spp. *and K. oxytoca* (see Table S 1).

### DNA extraction and Whole Genome Sequencing

The lysate from the bacterial colonies was obtained by suspending a single colony in 600 μl 1xPBS buffer with 1 mM EDTA and boiling for 10 minutes at 95°C. Genomic DNA for some isolates was extracted using DNeasy Blood & Tissue Kit (Qiagen), according to the manufacturer’s protocol. For other isolates, genomic DNA was obtained by automated purification using the MagNA Pure 96 DNA and Viral NA Small Volume Kit and DNA Blood ds SV 2.0 protocol (Roche Diagnostics). The DNA was quantified with Qubit fluorometer (Invitrogen, Waltham, MA, USA). DNA libraries were constructed using the Nextera XT library prep kit (Illumina, San Diego, USA) following the manufacturer’s protocol. Whole Genome Sequencing (WGS) was performed using Illumina NextSeq (2 × 150-bp paired-end sequencing). Quality control of the sequenced data was performed using Bifrost (https://github.com/ssi-dk/bifrost).

### Genomic characterization of the strains sequenced in this study

Assemblies were obtained using CLC Genomics (20). Species and Sequence-Types (STs) were identified using Kleborate 2.0.0 (21) (Table S 3).

## Data availability

The protocol used for Kp1 PCR target was designed and is available in protocols.io (dx.doi.org/10.17504/protocols.io.b4s5qwg6). Genomic sequences generated in this study were submitted to the European Nucleotide Archive and are accessible under the BioProject number PRJEB34643 and are also publicly available in BIGSdb (https://bigsdb.pasteur.fr/klebsiella/) through project ID 81 “Genomic dataset used for the validation of the triplex qPCR (KpSC + Kp1).”

---

**Figure S1.**
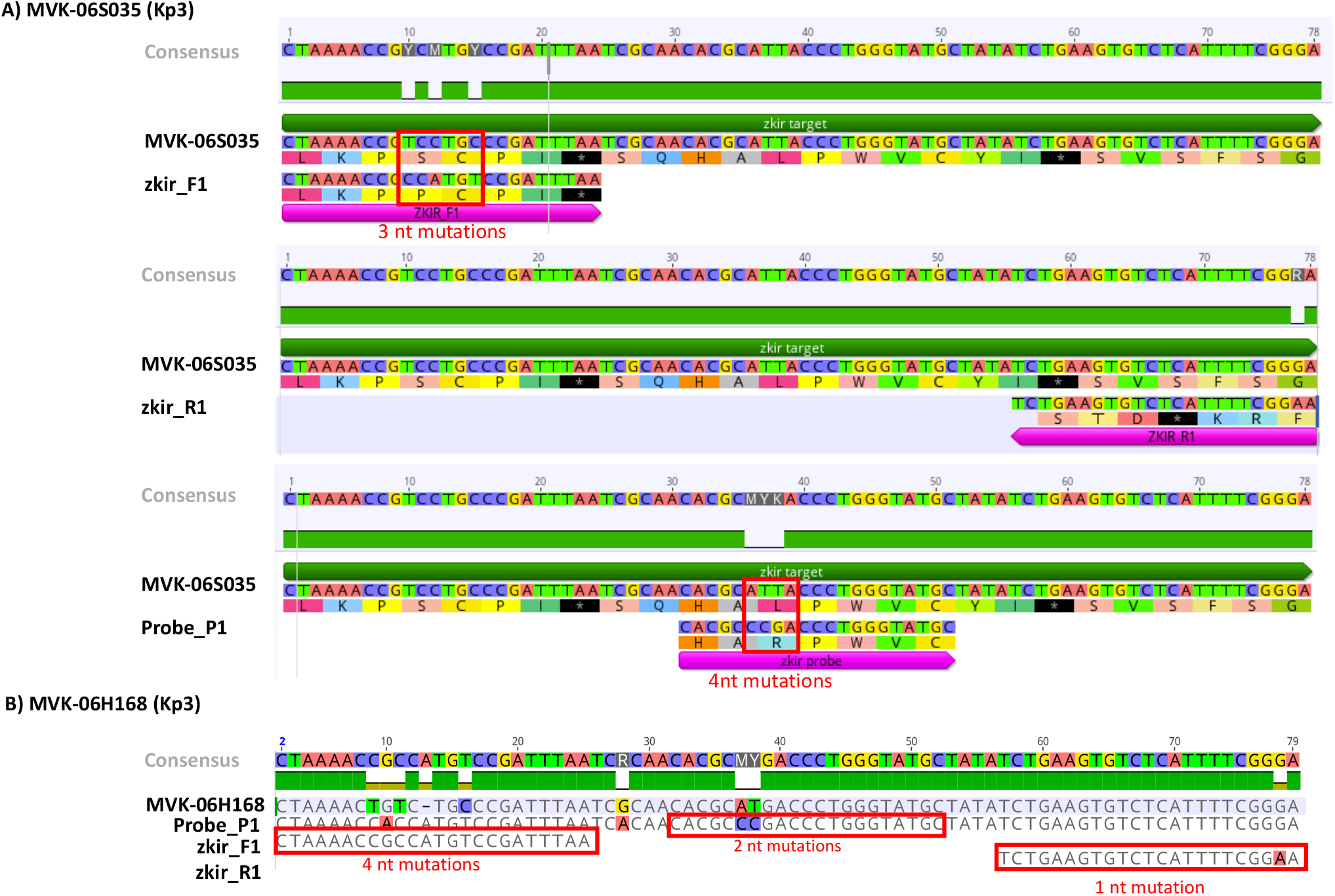
A) Mismatches in the forward primer and the probe region in MVK-06S035 strain (Kp3). B) Mismatches in the forward and reverse primer and the probe region in MVK-06H168 strain (Kp3).

