## Supplementary appendix for "Triplex real-time PCR ZKIR-T assay for simultaneous detection of the *Klebsiella pneumoniae* species complex and identification of *K. pneumoniae* sensu stricto"

### Supplementary material

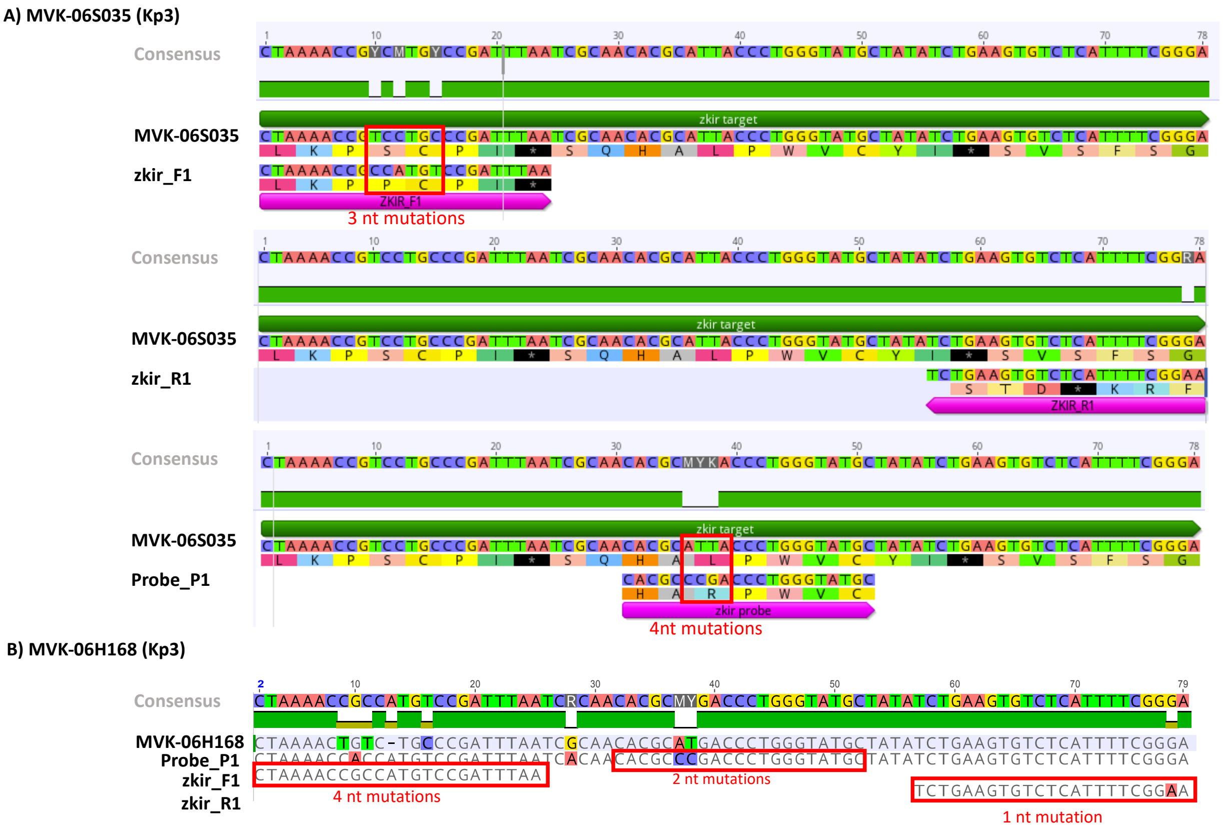


Figure S 1 A) Mismatches in the forward primer and the probe region in MVK-06S035 strain (Kp3). B) Mismatches in the forward and reverse primer and the probe region in MVK-06H168 strain (Kp3).

Table S 1. Bacterial strains used to develop the triplex assay (49 K. pneumoniae species complex (KpSc), 19 Klebsiella non-KpSC) at INRAe. Characterization and cycle threshold (Ct) values of the 49 KpSC strains representing phylogroups Kp1 to Kp7 and 19 closely related species.

| Species (Phylogroup) | Strain name | Strain bank ID* | Sampling date | Source | Ct_zkir_P1 | Ct_zkir_P2 | Ct_Kp1_P |
|---|---|---|---|---|---|---|---|
| ***Klebsiella pneumoniae* subsp. *pneumoniae* (Kp1)** | SB4-2 | SB1067 | 2002 | Feces | **17.6** | U | **23.5** |
| | ATCC13883$^T$ | SB132 | n.a. | Blood | **16.4** | U | **22.7** |
| | ATCC 700721 | SB107 | 1994 | Blood | **13.8** | U | **20.2** |
| | none | SB1139 | 2002 | Feces | **15.1** | U | **20.9** |
| | 5-2 | SB617 | 2000 | Natuurgebied canal | **15.6** | U | **21.3** |
| | MIAE07651 | none | 2015 | Blood | **17.3** | U | **23.2** |
| | 04A025 | SB20 | 1997 | Blood | **15.6** | U | **21.6** |
| | 2-3 | SB612 | 2000 | Rijnhauwen bridge | **15.5** | U | **21.6** |
| | BJ1-GA | SB4496 | 2011 | Liver abscess | **14.8** | U | **20.9** |
| ***Klebsiella quasipneumoniae* subsp. *quasipneumoniae* (Kp2)** | 01A030$^T$ | SB11 | 1997 | Blood | **19.5** | U | U |
| | none | SB1124 | 2002 | Canal water | **28.3** | U | U |
| | U41 | SB2110 | 1990 | Environment | **26.7** | U | U |
| | 10A442 | SB224 | 1998 | Blood | **18.5** | U | U |
| | 99-1002 | SB2478 | 1999 | n.a. | **18.3** | U | U |
| | 18A451 | SB255 | 1998 | Blood | **17** | U | U |
| | 11128 | SB3445 | n.a. | Diarrhoea | **20** | U | U |
| | CRBIP28.152 (18A69) | SB59 | 1997 | Blood | **26.1** | U | U |
| | KlebAli 0320584 | SB98 | n.a. | Environment | **18.8** | U | U |
| ***Klebsiella variicola* subsp. *variicola* (Kp3)** | 01A065 | SB1 | 1997 | Blood | **22.9** | U | U |
| | 07A058 | SB31 | 1997 | Blood | **21.6** | U | U |
| | IPEUC-1516 | SB3278 | 1988 | Urine | **22.9** | U | U |
| | CIP 53.24 | SB3295 | n.a. | n.a. | **22.5** | U | U |
| | CIP 53.26 (1756/51) | SB3301 | n.a. | n.a. | **23.3** | U | U |
| | F2R9$^T$ | SB48 | n.a. | Banana | **22.7** | U | U |

| | | | | | | | |
|---|---|---|---|---|---|---|---|
| | 6115 (KLSP49) | SB489 | n.a. | n.a. | **21.7** | U | U |
| | 4425/51 | SB497 | n.a. | n.a. | **21.8** | U | U |
| | Kp342 | SB579 | n.a. | Maize | **21.7** | U | U |
| ***Klebsiella quasipneumoniae* subsp. *similipneumoniae* (Kp4)** | CRBIP28.12 (09A323) | SB164 | 1997 | Blood | **26** | U | U |
| | 12A476 | SB203 | 1998 | Blood | **25.5** | U | U |
| | 07A044$^T$ | SB30 | 1997 | Blood | **25.6** | U | U |
| | 325 | SB3233 | 1975 | n.a. | **23.7** | U | U |
| | CIP 52.200 (1303/50) | SB3297 | n.a. | n.a. | **23.9** | U | U |
| | 1303/50 (KLSP50) | SB490 | n.a. | n.a. | **25.1** | U | U |
| | 4463/52 (KLSP60) | SB500 | n.a. | n.a. | **24.2** | U | U |
| | CIP110288 | SB4697 | 2010 | Farmland soil | **26.6** | U | U |
| | 1-1 | SB610 | 2000 | Lake kikker | **25.6** | U | U |
| ***Klebsiella variicola* subsp. *tropica* (Kp5)** | CDC 4241-71 | SB94 | n.a. | Environment | **22.4** | U | U |
| | Gal12 | SB824 | n.a. | Environment | **22.3** | U | U |
| | 814 | SB5387 | 2015 | Fecal sample | **20.3** | U | U |
| | 885 | SB5439 | 2016 | n.a. | **25.9** | U | U |
| | 1266$^T$ | SB5531 | 2016 | Fecal sample | **29.9** | U | U |
| | 1283 | SB5544 | 2016 | Fecal sample | **29.7** | U | U |
| | 1375 | SB5610 | 2016 | Fecal sample | **19.2** | U | U |
| ***Klebsiella quasivariicola* (Kp6)** | 08A119 | SB33 | 1997 | Blood | U | **27** | U |
| | 10982 | SB6071 | 2005 | Peri-rectal | U | **26.3** | U |
| | 01-467-2ECBU | SB6094 | 2015 | Feces | U | **26.1** | U |
| | 01-310A | SB6095 | 2013 | Vaginal swab | U | **27** | U |
| | KPN1705$^T$ | SB6096 | 2014 | Wound | U | **26.2** | U |
| ***Klebsiella africana* (Kp7)** | 200023$^T$ | SB5857 | 2016 | n.a. | **17.3** | U | U |
| ***Klebsiella michiganensis* (Ko1)** | CIP 110787$^T$ | SB4934 | 2010 | n.a. | U | U | U |
| | 05A071 | SB71 | 1997 | n.a. | U | U | U |
| | 09A029 | SB78 | 1997 | n.a. | U | U | U |
| ***Klebsiella grimontii* (Ko6)** | 07A479 | SB324 | 1998 | n.a. | U | U | U |
| | 06D090 | SB352 | 1998 | n.a. | U | U | U |

| | | | | | | | |
|---|---|---|---|---|---|---|---|
| | $06D021^{T}$ | SB73 | 1997 | n.a. | U | U | U |
| ***Klebsiella oxytoca* (Ko2)** | ATCC $13182^{T}$ | SB175 | n.a. | n.a. | U | U | U |
| | 02A067 | SB131 | 1997 | n.a. | U | U | U |
| | NCTC 49131 | SB136 | n.a. | n.a. | U | U | U |
| ***Klebsiella terrigena*$^{R}$** | $ATCC33257^{T}$ | SB170 | n.a. | n.a. | U | U | U |
| | 17C143 | SB313 | 1998 | n.a. | U | U | U |
| | V9813596 | SB2796 | 1998 | n.a. | U | U | U |
| ***Klebsiella planticola*$^{R}$** | 01A041 | SB7 | 1997 | n.a. | U | U | U |
| | $ATCC33531^{T}$ | SB174 | n.a. | n.a. | U | U | U |
| | 12C169 | SB303 | 1998 | n.a. | U | U | U |
| ***Klebsiella ornithinolytica*$^{R}$** | $ATCC31898^{T}$ | SB171 | n.a. | n.a. | U | U | U |
| ***Klebsiella aerogenes*** | CIP 60.86T | SB3629 | n.a. | n.a. | U | U | U |
| | 01A089 | SB538 | 1997 | n.a. | U | U | U |
| | 02A002 | SB539 | 1997 | n.a. | U | U | U |

U stands for undetermined, i.e. no amplification; Ct - cycle threshold, T indicates a type strain, n.a. - not available, R – also called *Raoultella* spp. according to (1)

Table S 2. Control strains representing each of the seven phylogroups from *K. pneumoniae* species complex used at SSI for the triplex PCR

| Strain name | Original Strain name | Phylogroup | Species |
|---|---|---|---|
| **MVK-06H079** | ATCC 13883$^T$ | Kp1 | *K. pneumoniae* |
| **MVK-06H171** | ATCC 700721 (MGH 78578) | Kp1 | *K. pneumoniae* |
| **MVK-06H172** | BIP28.152 | Kp2 | *K. quasipneumoniae* subsp. *quasipneumoniae* |
| **MVK-06H173** | CIP 53.26 (1756/51) | Kp3 | *K. variicola* subsp. *variicola* |
| **MVK-06H174** | CIP 52.200 (1303/50) | Kp4 | *K. quasipneumoniae* subsp. *similipneumoniae* |
| **MVK-06H175** | CDC 4241-71 | Kp5 | *K. variicola* subsp. *tropica* |
| **MVK-06H176** | 08A119 | Kp6 | *K. quasivariicola* |
| **MVK-06H177** | 200023$^T$ | Kp7 | *K. africana* |

$^T$ indicates a type strain.

Table S 3. Characterization and Ct values of 46 test strains representing phylogroups Kp1 to Kp6.

| Strain name | Sampling date | Source | ST | Species (PhG) [1] | Ct_zkir_P1 | | | Ct_zkir_P2 | | | Ct_Kp1_P | | |
|---|---|---|---|---|---|---|---|---|---|---|---|---|---|
| | | | | | Rep1 | Rep2 | Rep3 | Rep1 | Rep2 | Rep3 | Rep1 | Rep2 | Rep3 |
| **MVK-06S001** | feb-18 | Sewage | ST391 | *K. pneumoniae* (Kp1) | **18.2** | **17.8** | **18.1** | U | U | U | **20.9** | **21** | **21.3** |
| **MVK-06S005** | feb-18 | Sewage | ST391 | | **18.1** | **18.1** | **18** | U | U | U | **21.1** | **21.7** | **21.5** |
| **MVK-06S008** | feb-18 | Sewage | ST234 | | **17.6** | **17.8** | **18** | U | U | U | **20.6** | **21.4** | **21.4** |
| **MVK-06S009** | feb-18 | Sewage | ST976 | | **17.2** | **17.6** | **18.1** | U | U | U | **20.3** | **21.2** | **21.5** |
| **MVK-06S010** | feb-18 | Sewage | ST5765 | | **17.5** | **17.4** | **18.4** | U | U | U | **20.5** | **20.9** | **21.9** |
| **MVK-06S013** | feb-18 | Sewage | ST252 | | **17.6** | **17.6** | **18.5** | U | U | U | **20.6** | **20.9** | **21.6** |
| **MVK-06G003** | jun-18 | Animal carriage | ST5 | | **17.6** | **18.1** | **18.5** | U | U | U | **20.6** | **21.4** | **21.6** |
| **MVK-06G005** | jun-18 | Animal carriage | ST5 | | **17.3** | **17.8** | **18.1** | U | U | U | **20.2** | **21.3** | **21.3** |
| **MVK-06G009** | jun-18 | Animal carriage | ST661 | | **17.8** | **17.5** | **18.3** | U | U | U | **20.9** | **20.9** | **21.8** |
| **MVK-06G014** | jun-18 | Animal carriage | ST661 | | **17.7** | **17.6** | **18.3** | U | U | U | **20.8** | **21.1** | **21.7** |
| **MVK-06G017** | jun-18 | Animal carriage | ST46 | | **17.8** | **17.8** | **18.6** | U | U | U | **20.5** | **20.6** | **21.6** |
| **MVK-06G018** | jun-18 | Animal carriage | ST5 | | **17.8** | **17.6** | **18.4** | U | U | U | **20.5** | **20.9** | **21.5** |

| | | | | | | | | | | | | | |
|---|---|---|---|---|---|---|---|---|---|---|---|---|---|
| **MVK-06G021** | jun-18 | Animal carriage | ST661 | | **17.7** | **17.7** | **18.8** | U | U | U | **20.7** | **21** | **22.2** |
| **MVK-06G024** | jun-18 | Animal carriage | ST661-1LV[1] | | **17.9** | **18** | **18.4** | U | U | U | **21.1** | **21.5** | **21.8** |
| **MVK-06S016** | sep-18 | Sewage | ST391 | | **17.6** | **17.1** | **18.2** | U | U | U | **21.4** | **21.1** | **21.5** |
| **MVK-06S023** | sep-18 | Sewage | ST45 | | **18.4** | **17.4** | **18.2** | U | U | U | **21.7** | **20.9** | **21.5** |
| **MVK-06S029** | sep-18 | Sewage | ST391 | | **17.6** | **17.6** | **18.1** | U | U | U | **20.5** | **21** | **21.4** |
| **MVK-06S033** | sep-18 | Sewage | ST889 | | **17.5** | **17.8** | **18.1** | U | U | U | **20.6** | **21.2** | **21.6** |
| **MVK-06S038** | sep-18 | Sewage | ST273-2LV[2] | | **17.4** | **17.5** | **18** | U | U | U | **20.7** | **21.3** | **21.4** |
| **MVK-06S049** | sep-18 | Sewage | ST20-1LV[1] | | **17.4** | **17.8** | **17.9** | U | U | U | **20.8** | **21.7** | **21.2** |
| **MVK-06S045** | sep-18 | Sewage | ST4756 | *K. quasipneumoniae* subsp. *quasipneumoniae* (Kp2) | **28.6** | **27.7** | **29.2** | U | U | U | U | U | U |
| **MVK-06S012** | feb-18 | Sewage | ST5766 | *K. variicola* subsp. *variicola* (Kp3) | **25.3** | **24.5** | **25.5** | U | U | U | U | U | U |
| **MVK-06S024** | sep-18 | Sewage | ST5065 | | **25.1** | **24.4** | **26.5** | U | U | U | U | U | U |
| **MVK-06S035** | sep-18 | Sewage | ST285 | | **U** | **37.2** | **U** | U | U | U | U | U | U |
| **MVK-06S039** | sep-18 | Sewage | ST6110 | | **24.5** | **23.5** | **24.6** | U | U | U | U | U | 35.2 |
| **MVK-06H010** | apr-18 | Human clinical (feces) | ST3982 | | **25.8** | **25.4** | **26.3** | U | U | U | U | U | U |
| **MVK-06H054** | may-18 | Human clinical (feces) | ST641 | | **25.8** | **25.2** | **26.4** | U | U | U | U | U | U |
| **MVK-06H070** | may-18 | Human clinical (feces) | ST1562 | | **24.9** | **24.8** | **26.4** | U | U | U | U | U | U |
| **MVK-06H077** | jun-18 | Human clinical (feces) | ST1915 | *K. variicola* subsp. *variicola* (Kp3) | **25.3** | **24.7** | **25.8** | U | U | U | U | U | U |
| **MVK-06F135** | mar-19 | Food (salad) | ST146 | | **25** | **25.1** | **25.6** | U | U | U | U | U | U |
| **MVK-06F137** | mar-19 | Food (salad) | ST363 | | **25** | **31.4** | **25.6** | U | U | U | U | U | U |
| **MVK-06F149** | mar-19 | Food (salad) | ST662 | | **24.9** | **24.2** | **26.4** | U | U | U | U | U | U |
| **MVK-06H098** | may-19 | Human clinical (blood) | ST5771 | | **24.1** | **23.8** | **26.1** | U | U | U | U | U | U |
| **MVK-06H100** | may-19 | Human clinical (blood) | ST250 | | **24.5** | **23.8** | **25.9** | U | U | U | U | U | U |
| **MVK-06H113** | may-19 | Human clinical (blood) | ST355 | | **22.9** | **22** | **26.2** | U | U | U | U | U | U |
| **MVK-06H116** | may-19 | Human clinical (blood) | ST2594 | | **24.8** | **23.5** | **26.5** | U | U | U | U | U | U |
| **MVK-06H127** | may-19 | Human clinical (blood) | ST355 | | **24** | **23.5** | **25.8** | U | U | U | U | U | U |
| **MVK-06H156** | may-19 | Human clinical (blood) | ST5778 | | **24.2** | **23.9** | **25.9** | U | U | U | U | U | U |
| **MVK-06H166** | may-19 | Human clinical (blood) | ST5779 | | **24.4** | **24.1** | **25.7** | U | U | U | U | U | U |
| **MVK-06H167** | may-19 | Human clinical (blood) | ST208 | | **25.1** | **24.5** | **26.5** | U | U | U | U | U | U |
| **MVK-06H168** | may-19 | Human clinical (blood) | ST5798 | | **33.7** | **32.2** | **34.5** | U | U | U | U | 37.8 | U |
| **MVK-06F021** | sep-18 | Food (meat) | ST3450 | *K. quasipneumoniae* subsp. *similipneumoniae* (Kp4) | **36.1** | **35.3** | **37.4** | U | U | U | U | U | U |
| **MVK-06S018** | sep-18 | Sewage | ST367 | | **36.4** | **33** | **37** | U | U | U | U | U | U |
| **MVK-06H092** | may-19 | Human clinical (blood) | ST5770 | | **39.7** | **37.4** | **36.3** | U | U | U | U | U | U |

| | | | | | | | | | | | | | |
|---|---|---|---|---|---|---|---|---|---|---|---|---|---|
| **MVK-06H095** | may-19 | Human clinical (blood) | ST1822 | | **35** | **32.3** | **36.2** | U | U | U | U | U | U |
| **MVK-06H101** | may-19 | Human clinical (blood) | ST4973 | *K. quasivariicola* (Kp6) | U | U | U | **26.8** | **26.7** | **27.7** | U | U | U |
| **MVK-06H171** | | Control strain | | *K. pneumoniae* (Kp1) | **20.2** | **20.2** | **18.3** | U | U | U | **24** | **24** | **21.5** |
| **MVK-06H079** | | Control strain | | *K. pneumoniae* (Kp1) | **18.3** | **17.9** | **18.4** | U | U | U | **21** | **21.2** | **21** |
| **MVK-06H172** | | Control strain | | *K. quasipneumoniae* subsp. *quasipneumoniae* (Kp2) | **29.1** | **28.2** | **28.7** | U | U | U | U | U | U |
| **MVK-06H173** | | Control strain | | *K. variicola* subsp. *variicola* (Kp3) | **24.5** | **24** | **26** | U | U | U | U | U | U |
| **MVK-06H174** | | Control strain | | *K. quasipneumoniae* subsp. *similipneumoniae* (Kp4) | **32.6** | **31.4** | **35.1** | U | U | U | U | U | U |
| **MVK-06H175** | | Control strain | | *K. variicola* subsp. *tropica* (Kp5) | **24.2** | **24** | **25.6** | U | U | U | U | U | U |
| **MVK-06H176** | | Control strain | | *K. quasivariicola* (Kp6) | U | U | U | **27.3** | **27.4** | **28.2** | U | U | U |
| **MVK-06H177** | | Control strain | | *K. africana* (Kp7) | **21.1** | **21.1** | **22.5** | U | U | U | U | U | U |

Positive control strains representing all phylogroups are numbered sequentially from MVK-06H171 to MVK-06H177.

Rep1-3 denote replicates, U stands for Undetermined, i.e. no amplification; ST – Sequence Type, Ct - cycle threshold

The unexpected results are marked with red font and are addressed in the text.

[1]partial *phoE*; [2] partial *phoE* and *mdh*
